# Seedling microbiota engineering using bacterial synthetic community inoculation on seeds

**DOI:** 10.1101/2023.11.24.568582

**Authors:** Gontran Arnault, Coralie Marais, Anne Préveaux, Martial Briand, Anne-Sophie Poisson, Alain Sarniguet, Matthieu Barret, Marie Simonin

## Abstract

Synthetic Communities (SynComs) are being developed and tested to manipulate plant microbiota and improve plant health. To date, only few studies proposed the use of SynCom on seed despite its potential for plant microbiota engineering. We developed and presented a simple, reproducible and effective seedling microbiota engineering method using SynCom inoculation on seeds. The method was successful using a wide diversity of SynCom compositions and bacterial strains that are representative of the common bean seed microbiota. First, this method enables the modulation of seed microbiota composition and community size. Then, SynComs strongly outcompeted native seed and potting soil microbiota and contributed on average to 80% of the seedling microbiota. We showed that strain abundance on seed was a main driver of an effective seedling microbiota colonization. Also, selection was partly involved in seed and seedling colonization capacities since strains affiliated to Enterobacteriaceae and Erwiniaceae were good colonizers while Bacillaceae and Microbacteriaceae were poor colonizers. Additionally, the engineered seed microbiota modified the recruitment and assembly of seedling and rhizosphere microbiota through priority effects. This study shows that SynCom inoculation on seeds represents a promising approach to study plant microbiota assembly and its consequence on plant fitness.

## Introduction

Plants are associated with many microorganisms that can modulate their fitness (Vannier et al 2019; Arnault et al 2023). In this context, microbiota engineering is gaining attention as a potential way to improve plant disease management (Malacrino et al 2022; Li et al 2021), plant resilience (Schmitz et al 2022) and plant biomass (Liu et al 2022) in a more sustainable agricultural framework (Trivedi et al 2021). One way to modulate the composition of the plant microbiota is to assemble several cultured microorganisms (i.e. bacterial or fungal strains) in Synthetic Communities (SynComs). SynComs represent means to establish causality between microbiota composition and plant fitness (Vorholt et al 2017) and could be designed to improve plant health (Shayanthan et al 2022). To date, SynComs have been primarily applied on soil (Liu et al 2021; Baas et al 2016), rhizosphere (Li et al 2021; Marín et al 2021) and phyllosphere (Chen et al 2020).

Use of SynCom for microbiota engineering in agriculture is still facing some concerns including (i) stability of the SynCom over time (ii) capacity of SynCom members to colonize plant tissues and (iii) ability to compete with native communities (Rocca et al 2021). This latter phenomenon is called community coalescence (Rillig et al 2015; Rocca et al 2021). It is generally difficult to predict the outcome of a coalescence event since the resulting community depends on both neutral (e.g. dispersal, drift) and niche-based processes (e.g. host selection, biotic interactions). One important parameter that could influence the outcome of community coalescence is the mixing ratio of the two communities, i.e. mass effect (Shmida & Wilson 1985). For instance, Chen et al showed that SynCom concentration was a key factor that influenced both the SynCom stability in plant tissues and the microbial interactions within the SynCom (Chen et al 2020). Hence, SynCom inoculation represents a promising tool in agriculture but more research is needed to improve their efficiency and yield more predictable coalescence outcomes with native communities.

Plant microbiota engineering using SynCom inoculation on seed is gaining attention as a potential way to reduce the amount of inoculum needed (Rocha et al 2019). Indeed, seed could be a relevant vector to transmit beneficial microbiota to seedlings and change the trajectory of plant microbiota assembly (Debray et al 2021). Still, to date, only few studies have used SynComs on seed (Dos Santos et al 2021; Armanhi et al 2021; Kaur et al 2022, Simonin et al 2023), despite the potential use of seed coating to deliver beneficial microorganisms to crops (Rocha et al 2019). Additionally, seed microbiota has been shown to promote seedling growth and protection against fungal disease (Pal et al, 2022). Thus, seed-borne taxa represent an untapped microbial resource to improve plant protection and yield. Seeds harbor a specific microbiota: microbial biomass and richness are reduced compared to other plant microbiota compartments (Guo et al 2021) and are highly variable from one seed to another (Chesneau et al 2022). Moreover, several studies report that only a fraction of the seed microbiota is transmitted to the seedlings (Chesneau et al 2022; Rochefort et al., 2021; Walsh et al., 2021). In this sense, Walsh et al (2021) argue that seed inoculants may exhibit reduced effectiveness in highly diversified and populated soil due to mass effect. On the contrary, Moroenyane et al (2021) contend that the preeminence of the seed microbiota renders it a promising candidate for microbiota engineering. Thus, this raises new challenges and questions before a possible application of SynCom on seeds: (1) How effective is SynCom colonization of seeds and seedlings? (2) What are the relative contributions of mass effect and selection on SynCom colonization of both seeds and seedlings? (3) What is the impact of SynComs seed-inoculation on recruitment of environmental microbes (e.g. seed- and soil-borne taxa)?

To answer these questions, we developed a reproducible method of SynCom inoculation on common bean seeds (*Phaseolus vulgaris L.*). A set of three experiments was performed (Fig1-A). (Exp. 1) A first experiment investigated the impact of SynCom concentration and seed disinfection on seed colonization. (Exp. 2) Then, the importance of mass effect on seedling colonization was studied in a coalescence context with seed-borne and soil-borne taxa. (Exp. 3) Finally, the importance of strain identity was assessed through inoculation of 12 SynComs of variable species richness and composition.

**Figure 1.**
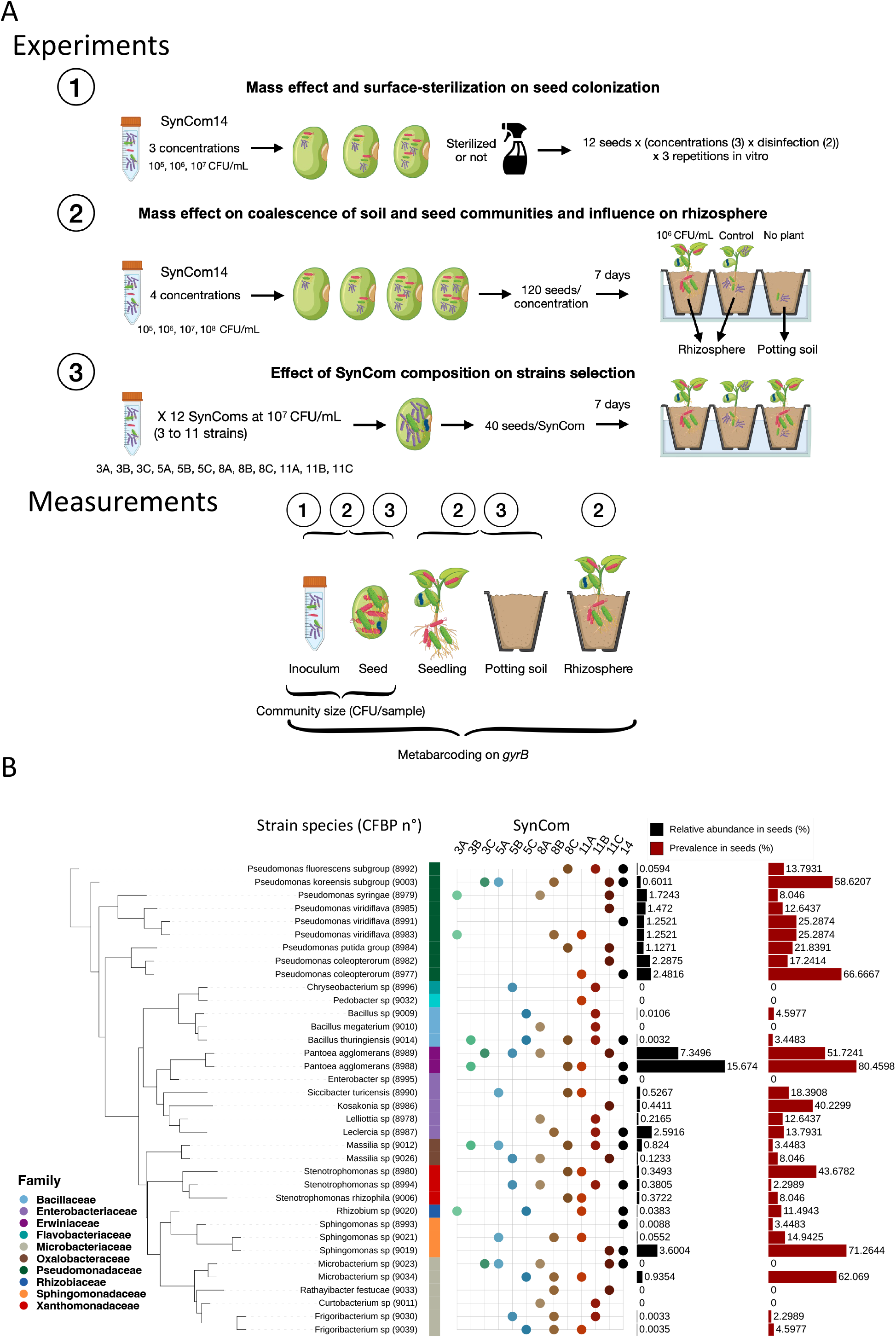
Design of the different experiments, strain selection and SynCom compositions. A) Overview of the different experiments. In Experiment 1, inoculation of SynCom14 (composed of 14 bacterial strains) on surface-sterilized and unsterilized seeds at different concentrations. In Experiment 2, influence of SynCom14 inoculation at different concentrations on seed and seedling microbiota assembly. In Experiment 3, influence of the inoculation of 12 different SynComs (with 3, 5, 8 or 11 bacterial strains) on seed and seedling microbiota assembly. B) Phylogenetic tree of the 36 strains selected and composition of the 13 SynComs. SynCom14 was studied in experiments 1 and 2 and the others in experiment 3. The number in SynCom names indicates the SynCom richness. Relative abundance and prevalence of each strain in the original seed samples are plotted on the right side. Seven strains were selected while they were not detected using the metabarcoding approach.

Since single seed harbored a microbiota of weak richness and biomass (Chesneau et al 2022) and that soil is the main inoculum source of seedling microbiota (Rochefort et al, 2021; Walsh et al, 2021) we hypothesize that (H1.1) potting soil will still be the main source of seedling microbiota. However, we expect that SynCom colonization will increase with the concentration applied through mass effects (H1.2). Also, we hypothesize that through selection processes (H2.1) SynCom strains will exhibit varying colonization capacities for both seeds and seedlings and that (H2.2) SynCom initial composition will influence strains transmission capacity through biotic interactions. Finally, through priority effects, we expect that inoculated seed will influence the recruited communities of both seedling (H3.1) and rhizosphere (H3.2).

## Materials and methods

### Plant material & Constitution of the culture-based collection of seed bacterial strains

Eight genotypes (Flavert, Linex, Facila, Contender, Vanilla, Deezer, Vezer, Caprice) of common bean (*Phaseolus vulgaris*) were grown on the field in 2020 by the National Federation of Seed Multiplicatiers (FNAMS) on two locations (Condom, Gers (43.956991, 0.392127) and Brain sur l’Authion, Maine-et-Loire (47.470532, -0.394526), FRANCE). These seed samples were used to obtain a collection of bacterial isolates from both seeds and seedlings. To isolate strains from seedlings germinated in gnotobiotic conditions, 30 seeds per condition were grown out in cotton soaked with 4mL of sterile water (growth during 7 days at 25°C, 16 hours of day, 8 hours of night) and then the seedlings were crushed and homogenized with 2mL of sterile water. The seeds were soaked in 2mL of sterilized water per gram of fresh material at 4°C under agitation (220rpm) overnight (∼16 hours). Suspensions were plated on tryptic soy agar 10% strength TSA and incubated at 18°C during 7 days. Isolated colonies were then picked up based on morphotype and grown in TSB in 96-well plates for 4 days at 18°C. A total of 1276 strains from these different seed and seedling samples were collected and stored at -80°C in 40% glycerol. The genotyping of the 1276 isolates was performed by metabarcoding of the *gyrB* bacterial marker gene (Illumina MiSeq). To assess the representativity of our strain collection, the total bacterial community composition of the seed (8 genotypes x 2 production sites x 3 replicates = 48 seed samples) and seedling (8 genotypes x 2 prod x 1 rep = 16 seedling samples) samples was characterized in parallel by metabarcoding of the *gyrB* marker following the protocol established by Barret et al. (2015) (see detailed method below).

From the 1276 isolates, 36 strains were selected (and deposited at the CIRM-CFBP strain collection) to recover variable relative abundance and prevalence on seed and seedling microbiota and covering a large phylogenetic diversity (FigS1, Fig1-B). DNA of these bacteria was extracted using the Wizard® Genomic DNA Purification Kit (Promega). Genomes were sequenced at the BGI (China) using DNBSEQ technology and assembled with Spades v3.15.3 using the default k-mer parameters (-k 21, 33, 55, 77, 99, 127) and the following options: -- cov-cutoff auto, --isolate (Prjibelski et al 2020). Genomes are available on NCBI using the BioProject ID PRJNA1041598.

### Design and inoculation of Synthetic Community on seeds

All experiments were made using a commercial seed lot of Flavert genotype from Vilmorin-Mikado (France). Strains and SynCom biomass of the inocula and the inoculated seeds biomass were verified using dilution and plating on 10% strength TSA. For each experiment, the control condition corresponds to the inoculation with sterile water. Control seeds were characterized using 3 seed batches of 25 seeds to get enough DNA amount for extraction.

#### 1. Mass effect and surface-sterilization on seed colonization (Exp. 1)

A first experiment was designed to adjust the SynCom inoculation protocol on bean seeds (experiment 1, Fig1-A). Especially, we tested the influence of inoculum concentration and seed surface disinfection on SynCom’s capacity to colonize seeds using a SynCom composed of 14 strains (hereafter SynCom14). Seeds were surface-sterilized using this protocol: sonicated for 1 min (40 Hz), soaked for 1 min in 96° ethanol, 5 min in 2.6% sodium hypochlorite, and 30 sec in 96° ethanol, and rinsed 3 times with sterile water. SynCom14 was designed to include a large taxonomic diversity of bacterial seed microbiota (Fig 1-B, FigS1) and was inoculated at different concentrations to study the impact of mass effect on seed colonization: 10^5^, 10^6^ and 10^7^ CFU/mL. To do so, each strain was resuspended in water by scratching a 48h culture of 10% strength TSA. Then, each strain was adjusted to an OD (600 nm) of ∼ 0.1 and SynCom was prepared by adding equivolume of each strain. Serial dilutions were made to obtain the final concentrations and strain and SynCom concentrations were checked using dilution and plating on 10% strength TSA. SynCom inoculations were performed by placing the seeds in a sterile container and adding 2mL of inoculum (sterile water for control) per gram of seed during 30 minutes under agitation (70 rpm) at 18°C. Excess inocula was then removed using a sterile strainer and seeds were dried during 30mn under laminar flow. For community size (CFU/seed), a total of 36 seeds per condition were studied (12 seeds per replicate x 3 independent experiment replicates) and for microbiota profiling a total of 24 seeds per condition were used (8 seeds x 3 replicates).

#### 2. Mass effect on coalescence of soil and seed communities and influence on rhizosphere microbiota (Exp. 2)

A second experiment aimed to study the relative importance of seed and potting soil communities during the establishment of seedling microbiota in a coalescence framework (experiment 2, Fig1-A). SynCom14 was inoculated as explained before at 4 different inoculum concentrations (10^5^, 10^6^, 10^7^ and 10^8^ CFU/mL) and 120 seeds per condition were germinated in non-sterile potting soil during 7 days in a growth chamber (16h day at 23°C, 8h night at 20°C, 70% humidity) to assess SynCom14 contribution to seedling microbiota. We also assessed the effects of SynCom inoculation on seeds on the rhizosphere microbiota. To do so, the potting soil bacterial communities were studied without seedling (no seedling), with 7 day seedlings that have not been inoculated (control seedling), and with seedlings inoculated with SynCom14 at 10^6^ CFU/mL. The adhering soil to seedling roots was considered as rhizosphere and conserved at -80°C before DNA extraction. Microbiota profiling was done on 8 seeds, 8 seedlings and 4 potting soil/rhizosphere per condition.

#### 3. Effect of SynCom composition on strain selection (Exp. 3)

A third experiment was designed to confirm the method using several SynCom compositions and of 4 levels of richness (3, 5, 8, 11 strains) to match the natural bacterial diversity observed on common bean mature seeds (Chesneau et al. 2022). For each given richness, strains were drawn randomly and without replacement using a pool of 33 strains (see Fig1-C and Fig S1 for SynComs composition and strain selection). A total of 40 seeds per condition were inoculated using a 10^7^ CFU/mL suspension following the same procedure described before and let grown in the same condition as described for experiment 2.

### Plant growth, DNA extraction and gyrB gene sequencing taxonomic classification

The following metabarcoding approach was performed on the inocula, inoculated seeds and seedlings of the different SynComs. For inocula, 200μL of each fresh inoculum was instantly stored at -80°C before DNA extraction. For inoculated seeds, individual seeds were soaked in 2mL of water overnight (∼16h) at 6°C under agitation (220rpm), 200μL of each suspension obtained was stored at -80°C before DNA extraction and suspension were plated to assess community size on seeds.

After 7 days of growth (BBCH stage 12, two full leaves unfolded), seedling roots were cleaned of potting soil excess by hand shaking and using sterilized water. Whole seedling samples were first crushed with a roller. Then 2mL of sterilized water was added and the samples were ground for 30s using a stomacher. DNA was extracted using 200μL of the crushed suspension with the NucleoSpin® 96 Food kit (Macherey-Nagel, Düren, Germany) following the manufacturer’s instructions. For potting soil and rhizosphere characterization, 4 replicates of 200mg per condition were extracted using DNA PowerSoil kit from Qiagen following the manufacturer’s instructions.

The first PCR was performed with the primers gyrB_aF64/gyrB_aR553 (Barret et al. 2015), which target a portion of *gyrB* gene in bacteria. PCR reactions were performed with a high-fidelity Taq DNA polymerase (AccuPrimeTM Taq DNA polymerase Polymerase System, Invitrogen, Carlsbad, California, USA) using 5µL of 10X Buffer, 1µL of forward and reverse primers (100µM), 0.2µL of Taq and 5 µl of DNA. PCR cycling conditions were done with an initial denaturation step at 94°C for 3 min, followed by 35 cycles of amplification at 94°C (30 s), 55°C (45 s) and 68°C (90 s), and a final elongation at 68°C for 10 min. Amplicons were purified with magnetic beads (Sera-MagTM, Merck, Kenilworth, New Jersey). The second PCR was conducted to incorporate Illumina adapters and barcodes. The PCR cycling conditions were: denaturation at 94°C (2 min), 12 cycles at 94°C (1 min), 55°C (1 min) and 68°C (1 min), and a final elongation at 68°C for 10 min. Amplicons were purified with magnetic beads and pooled. Concentration of the pool was measured with quantitative PCR (KAPA Library Quantification Kit, Roche, Basel, Switzerland). Amplicon libraries were mixed with 10% PhiX and sequenced with three MiSeq reagent kits v2 500 cycles (Illumina, San Diego, California, USA). A blank extraction kit control, a PCR-negative control and PCR-positive control (*Lactococcus piscium*, a fish pathogen that is not plant-associated) were included in each PCR plate. The raw amplicon sequencing data are available on the European Nucleotide Archive (ENA) with the accession number PRJEB59714.

The bioinformatic processing of the amplicons originating from the bacterial strain collection and SynCom experiments was performed in R. In brief, primer sequences were removed with cutadapt 2.7 (Martin, 2011) and trimmed fastq files were processed with DADA2 version 1.22.0 (Callahan et al., 2016). Chimeric sequences were identified and removed with the removeBimeraDenovo function of DADA2. Amplicon Sequence Variant (ASV) taxonomic affiliations were performed with a naive Bayesian classifier (Wang et al. 2007) with our in-house *gyrB* database (train_set_gyrB_v5.fa.gz) available upon request. Unassigned sequences at the phylum level and *parE* sequences (a *gyrB* paralog) were filtered. Then, only ASVs with a minimum of 20 reads and present in at least 2 different samples were retained for experiment 2 and 3 (3 reads in at least 3 samples for experiment 1). To track our SynCom strains, only ASVs with 100% of identity were considered to be our strains. Some single-nucleotide polymorphisms (SNPs) were identified for some strains and were artificially increasing inoculum richness. They were thus removed before downstream analyses. All R scripts employed in this work are available on GitHub (https://github.com/GontranArnault/BeanSeedSynCom2023).

### Statistical analyses and microbiota analysis

Microbial community analyses were conducted using the Phyloseq package v1.44.0 (Mc Murdie and Holmes 2013) using R. Figures were visualized using ggplot2 v3.4.3. Alpha diversity analyses were performed with a coverage-based rarefaction as recommended by Chao and Jost (2012) using iNEXT package v2.6.4 (Hsieh et al 2016). Beta diversity analyses were made using Bray-Curtis distance and permutational multivariate analysis of variance (adonis2 function of vegan v2.6.4 (Oksanen et al 2007), 999 permutations) after a rarefaction at 10000 reads per sample for experiments 1 and 3 and 6000 reads for experiments 2 (see rarefaction curves in FigS2). Beta diversity was visualized using Principal Coordinate Analysis (PCoA). For figure 6A, ASVs of the SynCom strains were removed to visualize the recruited communities’ assembly. For mean comparisons, ANOVA followed by post-hoc Tukey tests were conducted when the conditions of application were met. Otherwise, Kruskal-Wallis followed by pairwise Wilcoxon tests were conducted. P-values were corrected using Benjamini-Hochberg method when multiple comparisons were conducted.

To assess the relative contribution of native seed microbiota, potting soil and SynComs, a microbial source tracking analysis was conducted using FEAST v0.1.0 with a maximum iteration of 1000 (Shenhav, L. et al 2019). Control seed, inoculated seed and potting soil microbiota were considered as sources of microorganisms and seedling were considered as sink. The phylosignal package v1.3 (Keck et al 2016) was used to confirm the observed phylogenetic pattern between strain families and their colonization capacity of both seed and seedling. To do so, a phylogenetic tree was constructed using automlst (commit b116031) with default parameters and 1000 bootstrap replicates (-bs 1000) (Alanjary et al 2019). The local Moran’s index was calculated using the lipaMoran function. This index allowed us to test the positive or negative autocorrelations between the measured parameters (relative abundance on seed, seedling and their ratio) and the phylogenetic position of a given strain. Figure 1B was assembled using iTol. Changes in the relative abundance of bacterial communities of rhizospheres from experiment 2 were assessed with CornCob package v0.3.2 using taxatree_models function (log2 transformation) and taxatree_plots function for visualization.

## Results

### Seed colonization by SynCom depends on both mass effect and SynCom composition but not seed disinfection

A first SynCom, composed of 14 bacterial strains representative of the taxonomic diversity of our strain collection (SynCom14), was inoculated at three different concentrations on either surface sterilized or unsterilized bean seeds (Fig1, Exp1). Seed disinfection did not influence the number of CFU on seed after SynCom14 inoculation (Fig2-A). Bacterial richness (number of ASVs) was similar between native and disinfected seeds except for the lowest inoculated SynCom14 concentration (10^5^ CFU/mL) where disinfection reduced the number of ASVs (Fig2-B). Finally, seed community composition was significantly impacted by SynCom concentrations (R^2^ = 45.7%, p-value < 0.001), while seed disinfection was not driving changes in community composition (Fig2-C).

**Figure 2:**
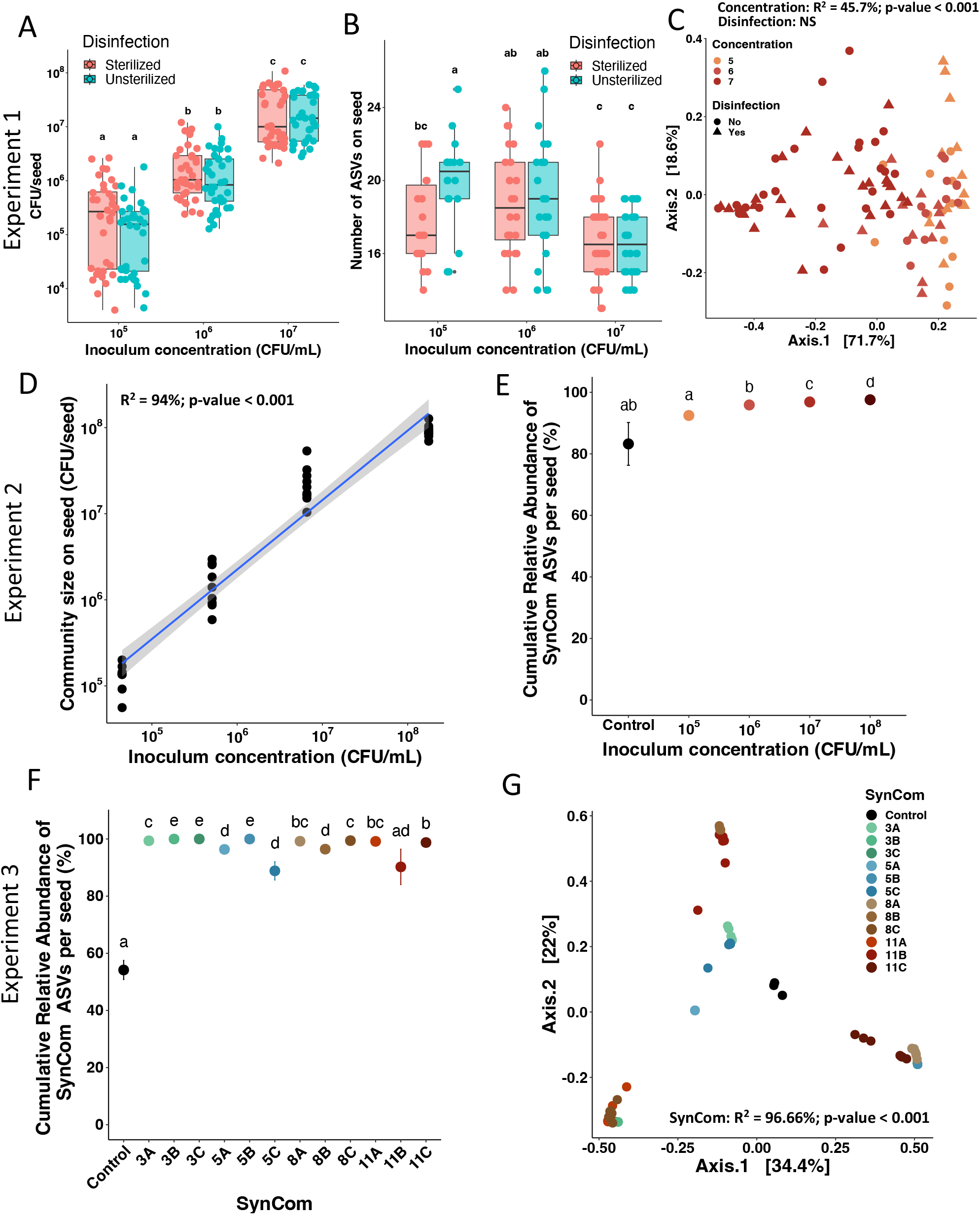
Effects of seed sterilization, SynCom concentration and composition on seed colonization. A) Community size on seed (CFU/seed) of SynCom14 in function of inoculum concentration (CFU/mL) and seed disinfection for the experiment 1. B) Number of ASVs detected on seeds inoculated with SynCom14 in experiment 1 depending on inoculum concentration and seed disinfection. C) Influence of SynCom14 concentration and seed disinfection on seed bacterial community structure visualized through a PCoA ordination based on Bray-Curtis distances (PERMANOVA test; Disinfection effect: non-significant, Concentration: R2 = 45.7%, p-value < 0.001). D) Correlation between community size on seed (CFU/seed) of SynCom14 in function of inoculum concentration (CFU/mL) for the experiment 2 (R2 = 94%, p-value < 0.001). E) Cumulative relative abundance of SynCom14 ASVs (SynCom colonization) in seeds from experiment 2 depending on inoculum concentration. F) Cumulative relative abundance of SynComs ASVs from experiment 3 (SynCom colonization) in seeds. G) Influence of SynCom composition of experiment 3 on seed bacterial community structure visualized through a PCoA ordination based on Bray-Curtis distances (PERMANOVA; SynCom condition: R2 = 96.66%, p-value < 0.001).

In a second independent experiment with non-sterilized seeds, changes in community sizes were correlated (R^2^ = 94%, p-value < 0.001) with the initial SynCom14 concentrations (Fig2-D). SynCom strains’ ASVs were detected on control seed samples (83% of relative abundance), as they are members of the native bean seed microbiota (Fig2-E and Fig4). The ASVs corresponding to the strains assembled in the SynCom14 represented on average 96% of the cumulative abundance of bacterial taxa detected on seeds, ranging from 92% at the lowest concentration (10^5^ CFU/mL) to 98% at the highest concentration (10^8^ CFU/mL) (Fig2-E).

In a third experiment, 12 SynComs were designed with four gradual levels of richness (3, 5, 8, 11 strains). ASVs of the corresponding strains ranged from 89% (SynCom 5C) to 99.9% of the seed relative abundance (SynCom 3C, Fig2-F). As expected, seed community composition was significantly explained by the inoculated SynCom inoculation (R^2^ = 96.7%, p-value < 0.001) (Figure 2-G).

Overall, these results showed that seed microbiota compositions were deeply modified by the SynCom inoculation and that SynCom concentration and composition were the main drivers of the overall seed composition.

### Seedling colonization by SynComs is driven by mass effects and initial SynCom composition

To find out whether the compositional changes observed in the seed persist during emergence, seedling microbiota were characterized 7 days post SynCom inoculation. The level of seedling colonization was estimated by monitoring the cumulative relative abundance of ASVs corresponding to strains assembled in the SynComs. ASVs of SynCom14 were detected on control seedlings at a low level (< 1%). In inoculated condition, ASVs of SynCom14 inoculation represented on average 87% of the seedling relative abundance, ranging from 75 to 93% depending on inoculum concentration (Fig3-A). A significant increase in ASVs of SynCom14 was observed between control, 10^5^ and 10^6^ CFU/mL before reaching a plateau (Fig3-A). Hence seedling bacterial composition was successfully modified by SynCom inoculation. According to beta-dispersion (distance to centroid) the variability in seedling community structure was significantly reduced following SynCom14 inoculation compared to non-inoculated seeds (Fig3-C). Moreover, seedling community composition was significantly influenced (R^2^ = 11.65%, p-value < 0.001) by SynCom14 concentration (Fig3-E). To assess whether SynCom richness could modify seedling microbiota composition, 12 SynComs of increasing strain richness were seed-inoculated at the same initial concentration (10^7^ CFU/mL). On average, SynComs ASVs represented 80% of seedling microbiota (Fig3-B). While SynComs colonizations of seedlings were more variable (29% for SynCom 5C to 95% for SynCom 8C), a very low influence of initial SynCom richness was detected (R^2^ = 12.7%, p-value < 0.001, FigS3). In contrast the initial SynCom composition resulted in different seedling colonization. The most prominent example concerned the three SynComs composed of 5 strains with approximately 80% of cumulative relative abundance for SynComs 5A and 5B and ∼30% for SynCom 5C (Fig3-B). Interestingly, these differences in seedling colonization were correlated with the initial SynCom establishment on seeds (Fig3-D, R^2^ = 44.94%, p-value < 0.001). Thus, one important parameter that could predict bacterial colonization of seedling was its community size on seed. Seedlings inoculated with the same SynCom clustered together and the PERMANOVA confirmed that 79.84% (p-value < 0.001) of the variance was explained by the SynCom inoculation (Fig3-D).

**Figure 3:**
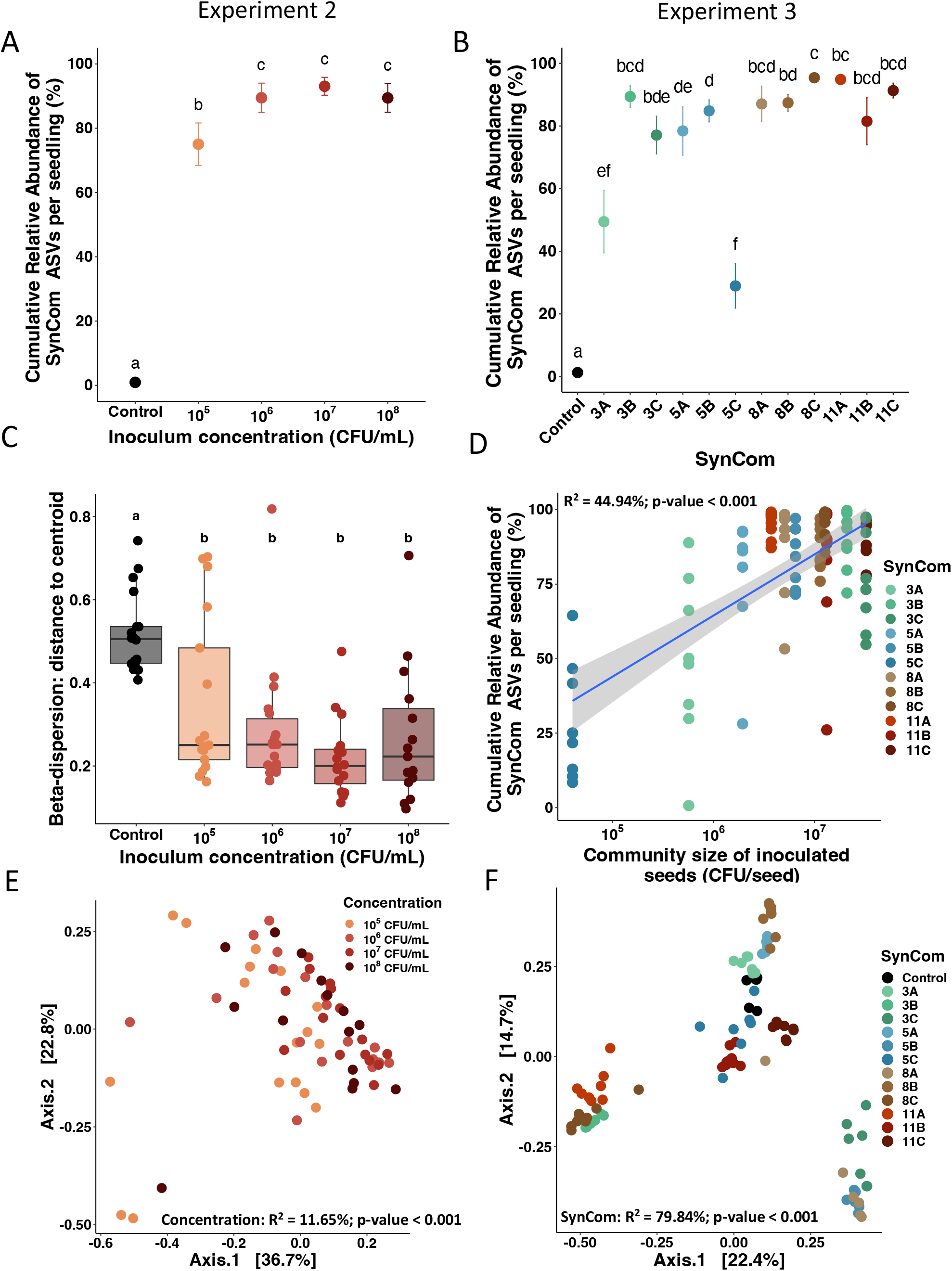
influence of SynCom mass effect and composition on seedling colonization. A-B) Cumulative relative abundance of SynComs ASVs (SynCom colonization) in seedlings from experiment 2 (A) and 3 (B). C) Beta-dispersion analysis (distance to centroid) of seedlings inoculated with SynCom14 (experiment 2) (Tukey test, the different letters indicate statistically different groups). D) Correlation between the cumulative relative abundance of SynCom ASVs in seedlings and the mean community size of inoculated seeds from experiment 3 (Pearson correlation R^2^ = 44.94, p-value < 0.001). E) Influence of SynCom14 concentration on seedling bacterial community structure visualized through a PCoA ordination based on Bray-Curtis distances (PERMANOVA; concentration: R^2^ = 11.65%, p-value < 0.001). F) Influence of SynCom composition from experiment 3 on seedling bacterial community structure visualized through a PCoA ordination based on Bray-Curtis distances. (PERMANOVA; concentration: R^2^ = 79.84%, p-value < 0.001).

Overall, these results showed that seedling microbiota of common bean could be deeply modified using SynCom inoculation on seeds and that this inoculation procedure greatly modified community composition observed on seedlings.

### Strains show heterogeneous transmission capacities from seed to seedling

Based on the unique *gyrB* ASV of each inoculated strain, we tracked the strain transmission from the inoculum to the seedling of experiment 3 (Fig4). All the strains’ ASVs were detected in at least 2 samples and were thus kept during the filtering process, except for the *Pedobacter sp* (CFBP9032) strain.

**Figure 4:**
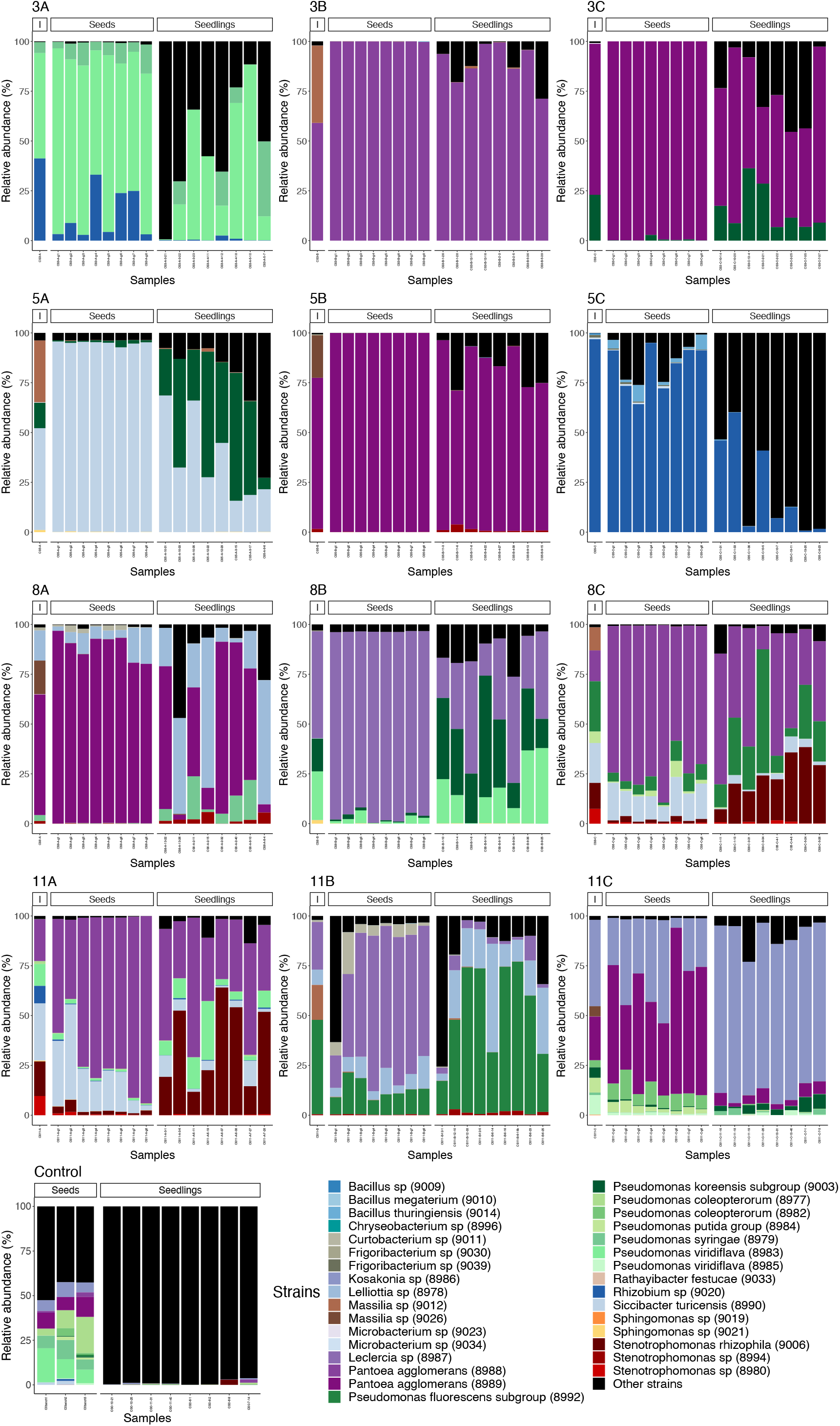
Influence of SynCom composition on taxonomic profiles of inocula, seeds and seedlings from experiment 3. Relative abundance of inoculated strains in the inocula, seeds and seedlings. Each stacked bar represents a sample. Only ASVs of the SynCom strains are colored, the black part represents uninoculated environmental taxa (e.g. potting soil & native seed microbiota). Per SynCom condition, 1 inoculum, 8 inoculated seeds and 8 seedlings were characterized using amplicon sequencing of the *gyrB* gene. For the control condition (not inoculated), 3 seed batches of 25 seeds and 8 individual seedlings were characterized.

The different SynCom panels confirmed the observation made on the overall community structures: seed and seedlings inoculated with SynComs showed very different taxonomic profiles depending on the inoculated SynCom (Fig4). In particular, each SynCom condition showed very distinct taxonomic profiles even at strain level. Also, taxonomic compositions of the SynComs were distinct between the inoculum, the seeds and the seedlings, highlighting the variability in strains’ ability to colonize the different habitats. For instance, in SynComs 8C and 11A, *Stenotrophomonas rhizophila* (CFBP9006) had a reduced relative abundance from inoculum to seeds but then increased from seeds to seedlings. For a given strain in each SynCom, the transmission rate from seed to seedlings was assessed based on presence of ASV in each habitat (Fig5-A). We identified 16 strains that had a transmission rate to seedlings of 100% in each SynCom tested. These strains are from the genera *Kosakonia*, *Leclercia*, *Pantoea*, *Pseudomonas*, *Rhizobium*, *Siccibacter*and *Stenotrophomonas*. On the other hand, *Microbacterium sp* (CFBP9034) and *Bacillus sp* (CFBP9009) strains had a transmission success of 0%. Some intermediate strains had interesting patterns: their transmission success was variable depending on the SynCom tested. For instance, *Sphingomonas sp* (CFBP9021) had a transmission rate of 75% in SynCom 8B and SynCom 5A and had a transmission rate of 0% in SynCom 11A. This example highlights the importance of strain interactions during seedling microbiota assembly (Fig5-A).

**Figure 5:**
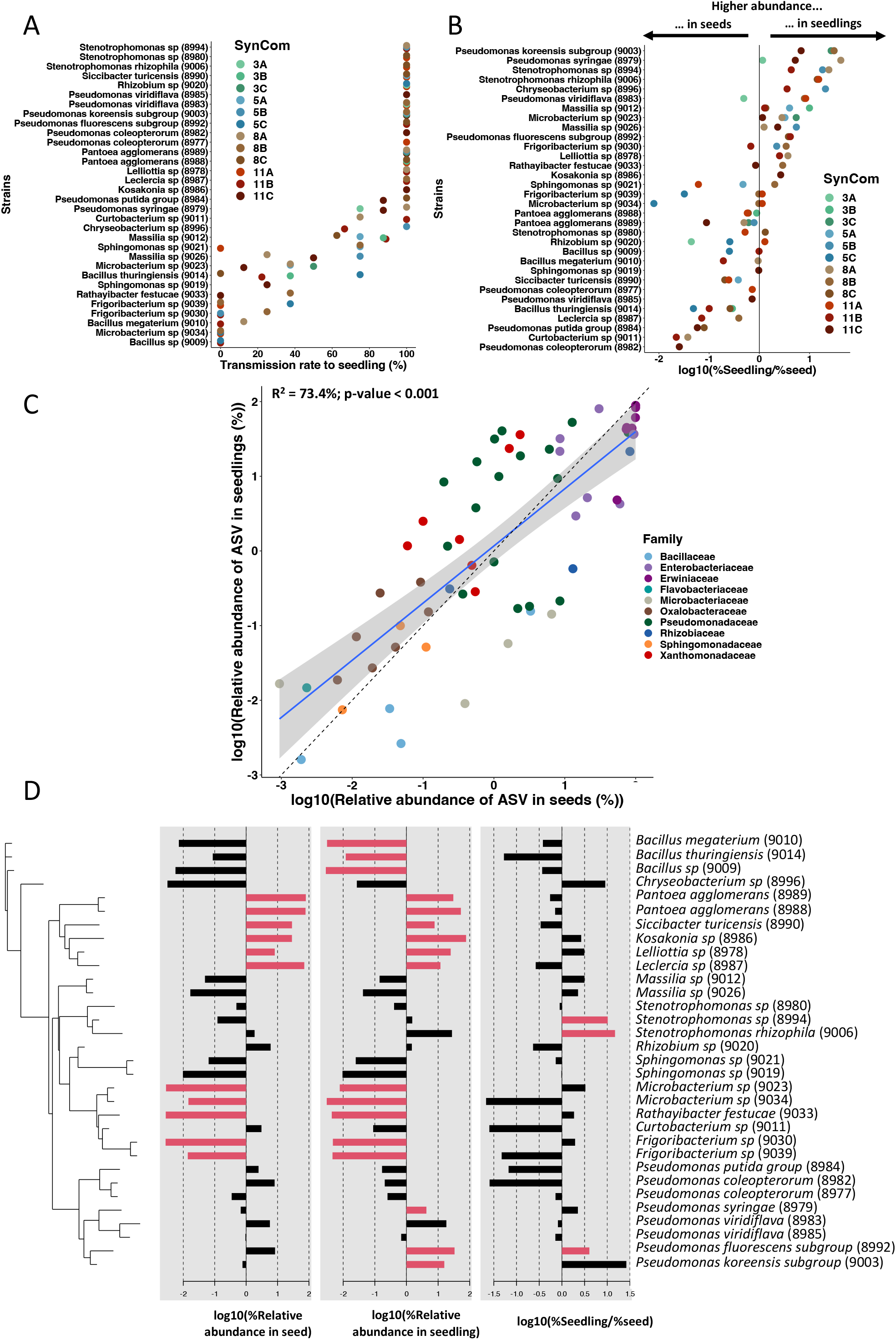
Transmission of each strain from seed to seedling. A) Transmission rate to the seedlings of each strain in the different SynComs of experiment 3. B) Strain’s ability to colonize seedlings compared to their initial relative abundance on seeds, depending on the SynCom composition of experiment 3. The following ratio was calculated to assess this trait: log10(Relative abundance on seedling/Relative abundance on seed). C) Correlation between each ASV relative abundance on seeds compared to their relative abundance on seedlings. A linear model confirms the relationship with this given equation: log10(%Seedling) = -0.02806+0.80771 x log10(%Seed) (R^2^ = 73.4%, p-value < 0.001). y = x dashed line was plotted to see if an ASVs is enriched or depleted in seedlings compared to its relative abundance on seeds (above or under the y = x dashed line respectively). D) Phylogenetic pattern of strains colonization ability to colonize seeds and seedlings. Phylosignal package was used to test the significance of the observed phylogenetic signal. The red barplots show the significantly (p-value < 0.05) enriched or depleted strains based on local Moran’s index, or Local Indicators of Phylogenetic Association (LISA), using the lipaMoran function.

To go further, we assessed the ability of each strain to colonize seedlings compared to their initial relative abundance on seeds, depending on the SynComs, using this ratio: log10(% Relative abundance on seedling / % Relative abundance on seed) (Fig5-B). Some strains were always found to be better seedling colonizers, including *Pseudomonas koreensis subgroup* (CFBP9003), *Pseudomonas syringae* (CFBP8979), *Stenotrophomonas sp* (CFBP8994), *S. rhizophila* (CFBP9006), *Chryseobacterium sp* (CFBP8996), *Massilia sp* (CFBP9012) and *Pseudomonas fluorescens subgroup* (CFBP8992). On the other hand, some strains were found to be better seed colonizers, including *Pseudomonas coleopterorum* (CFBP8982), *Curtobacterium sp* (CFBP9011), *Pseudomonas putida group* (CFBP8984), *Leclercia sp* (CFBP8987), *Bacillus thuringiensis* (CFBP9014), *Pseudomonas viridiflava* (CFBP8985), *Pseudomonas coleopterorum* (CFBP8977) and *Siccibacter turicensis* (CFBP8990). Finally, some strains had variable behaviors depending on the SynCom composition. For instance, the *Rhizobium sp* (CFBP9020) was a better seed colonizer in the SynCom 3A and 5C but a better seedling colonizer in the SynCom 11A.

Additionally, we found that the seedling relative abundance of an ASV in a given SynCom was correlated with its relative abundance in seed (Fig5-C, R^2^ = 73.4%, p-value < 0.001). Thus, the seedling relative abundance of a given ASV could be predicted based on its relative abundance on seeds. This confirms at the strain level that high seed colonization leads to high seedling colonization. A phylogenetic pattern was observed in this correlation: Enterobacteriaceae and Erwiniaceae (purple) were highly abundant on both seed and seedling. Bacillaceae (blue) and Microbacteriaceae (grey) were depleted in seed and seedling microbiota (below the y = x dashed line), while *Stenotrophomonas* sp (CFBP8994) and *S. rhizophila* (CFBP9006) were enriched in seedling microbiota (red, Xandomonadaceae, above the y = x dashed line). This phylogenetic pattern was confirmed statistically (Fig5-D).

To conclude, we confirm that the relative abundance of a strain on seeds is a good predictor of its future relative abundance on seedling. However, strains showed different capacities of seed and seedling colonization depending on their phylogeny.

### SynCom inoculation influences seedling and rhizosphere microbiota recruitment from environmental sources

The relative contribution of the different inoculum sources characterized (i.e. potting soil, native seed microbiota and inoculated seed microbiota, Fig6-A) was assessed with FEAST. Overall, the SynCom inoculation significantly decreased the unknown source proportion compared to control seedlings (Fig6-A and FigS4). The native seed microbiota had a significantly increased contribution in SynComs 3A and 5C and a significantly decreased contribution in SynComs 3B, 5A, 8C, 11B and 11C (Fig6-A and FigS4). The potting soil contribution was on average 4.9% (ranging from 0.9% in SynCom 8A to 12.8% in SynCom 5C) and was not significantly different between all the conditions (FigS4). Overall, SynCom composition influenced the contribution of the different microbial sources driving seedling microbiota assembly.

**Figure 6:**
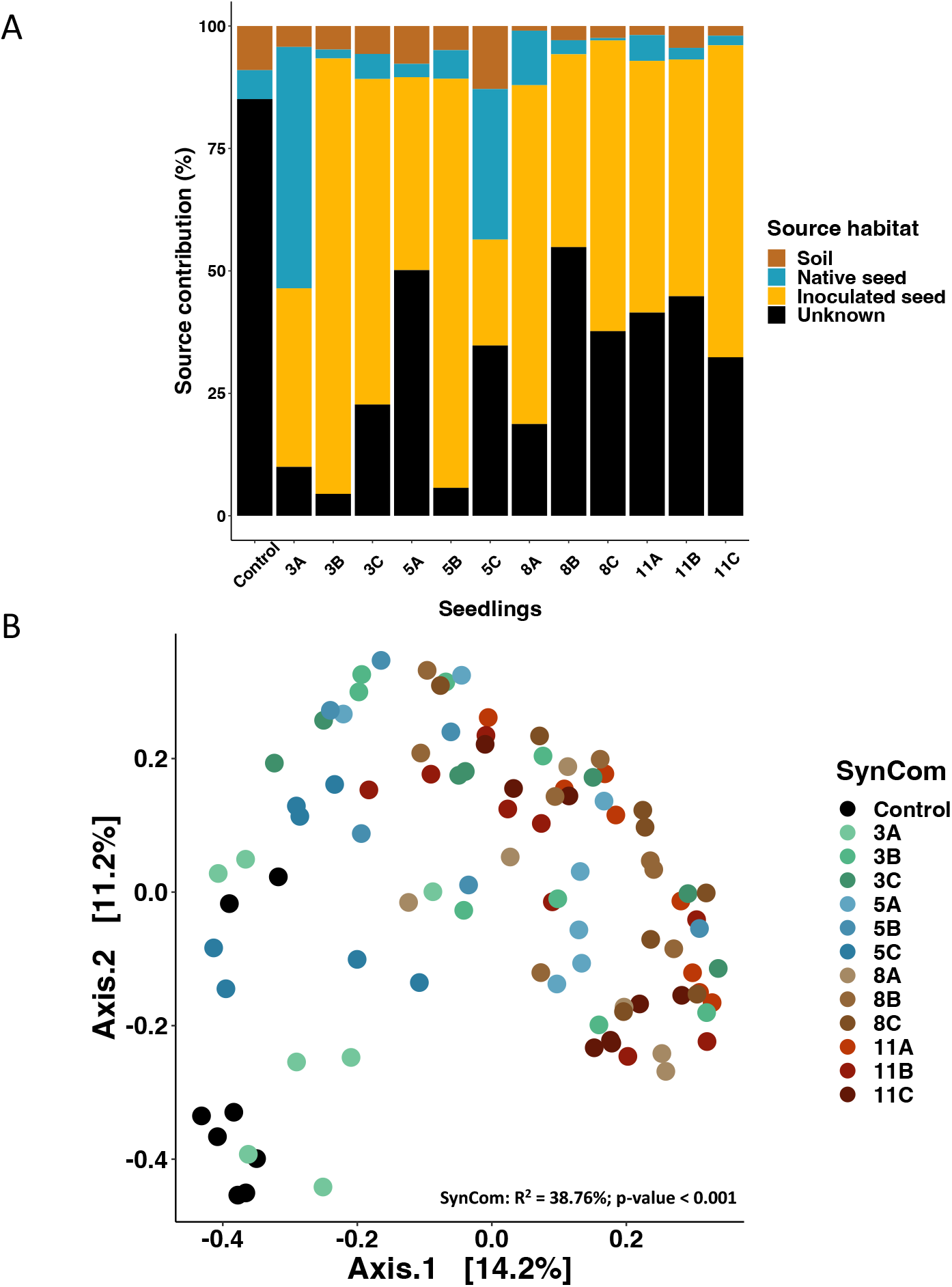
SynCom effect on environmental taxa recruitment of seedlings. A) To assess the relative contribution of native seed microbiota, potting soil and inoculated seed, a microbial source tracking analysis was conducted using FEAST. Control seed, inoculated seed and potting soil microbiota were considered as sources of microorganisms and seedling were considered as sink. (See detailed boxplot for each source and SynCom and associated statistics in figure S4). B) Influence of SynComs on seedling bacterial communities recruited from other sources (potting soil, air, water) visualized through a PCoA ordination based on Bray-Curtis distances (PERMANOVA; SynCom: R^2^= 38.76%, p-value < 0.001).

**Figure 7:**
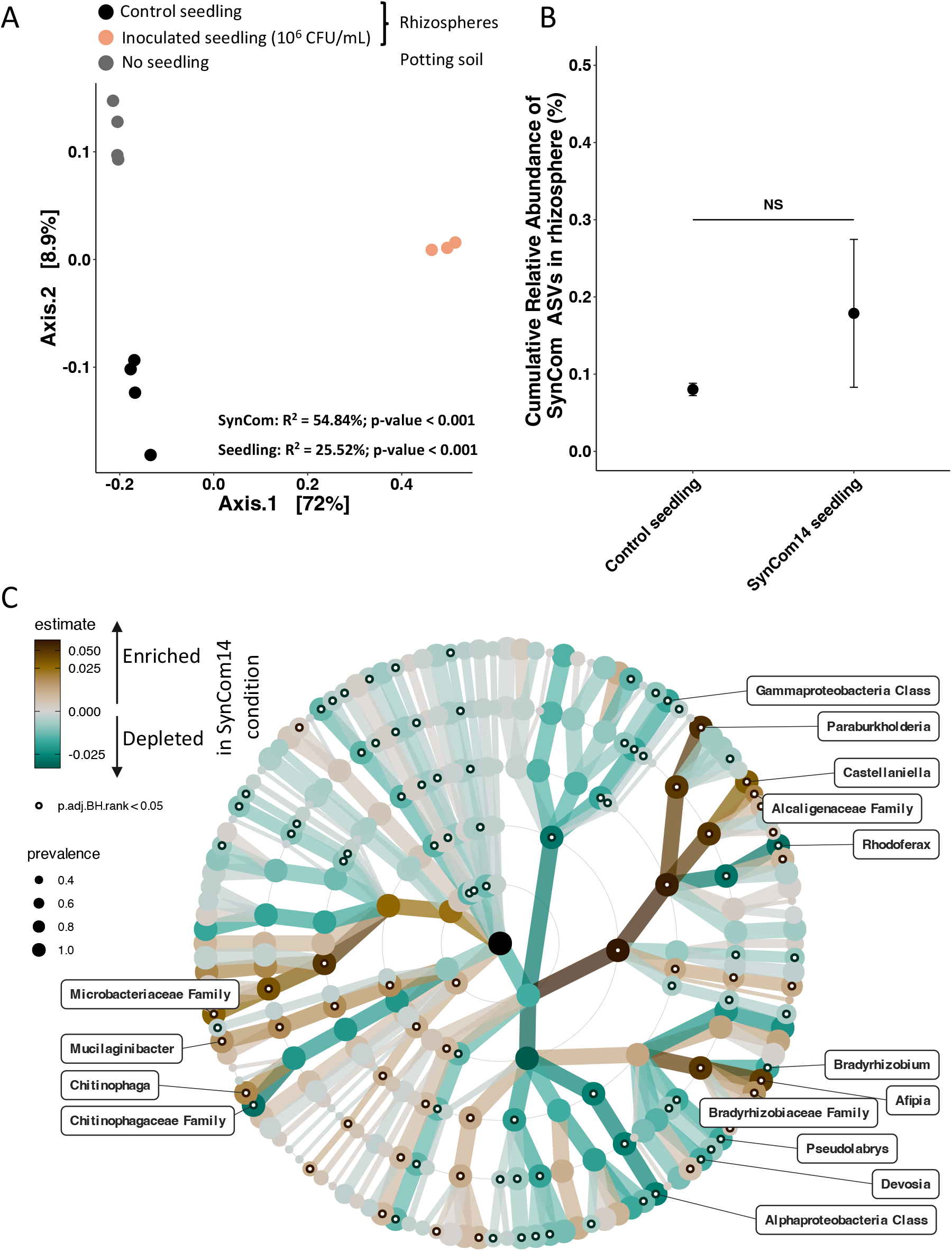
Effect of SynCom14 on rhizosphere community. A) Potting soil bacterial community structure visualized through a PCoA ordination based on Bray-Curtis distances. The potting soil bacterial communities were studied without seedling (No seedling), with a seedling from a non-inoculated seed (Control seedling), and with a seedling coming from inoculated-seed with the SynCom14 (PERMANOVA; SynCom: R^2^ = 54.84%, p-value < 0.001; Plant effect: R^2^ = 25.52%, p-value < 0.001). B) Cumulative relative abundance of SynCom14 ASVs in rhizospheres of control and inoculated plants. C) Changes in the relative abundance of bacterial genera of rhizospheres of inoculated plants (SynCom14) or in control plants (not inoculated) at the different taxa levels. Labels of the corresponding genera were plotted only if adjusted p-value < 0.05 and if estimate was below and above 0.01.

Although seedling microbiota was mainly composed of SynCom strains, about 20% of taxa were derived from other environmental sources. This provides an opportunity to assess the role of SynCom composition on recruitment of these taxa. After removing ASVs of SynComs strains, the similarities between seedling communities were assessed with Bray-Curtis distances (Fig6-B). Even in the absence of SynCom members, the inoculation of SynComs remained an important driver of seedling community structure (R^2^ = 38.76%, p-value = 0.001, Fig6-B). Taxonomic profile of the recruited microbiota was described but showed no clear pattern (FigS5-A). In seedlings derived from uninoculated seeds (control), one ASV of *Enterobacter cloacae* was highly abundant (56% of seedling microbiota, FigS5-B). The relative abundance of this ASV decreased in seedling from seeds inoculated with SynComs ranging from 0.012% in SynCom 8C to 16% in SynCom 3A (FigS5-B).

Next, we assessed the effects of SynCom inoculation on seeds on the rhizosphere microbiota. The potting soil bacterial communities were studied on day 7 without plant (No Plant), with a seedling that had not been inoculated (control plant), and with a seedling inoculated with one SynCom (SynCom14, Exp 2). Permanova analysis showed that 55% of the variance was explained by the inoculation while 26% was explained by the seedling presence itself (Fig7-A). ASVs corresponding to SynCom14 members had a cumulative relative abundance of <0.2% in the rhizospheres of control and inoculated plants (Fig7-B). Hence, SynCom14 members did not colonize and/or persist in the surrounding soil 7 days post-inoculation. Nevertheless, the inoculation of SynCom14 led to significant differences in taxonomic composition of the rhizospheres (Fig7-C). For instance, inoculation of SynCom14 on seeds led to an increased relative abundance of *Paraburkholderia*, *Castellaniella* and *Chitinophaga* and a decreased relative abundance of *Devosia, Rhodoferax and Pseudolabrys* compared to the control condition (Fig7-C). Thus, SynCom14 inoculated on seed at day 0 induced deep modifications on seedling rhizosphere composition at day 7 without colonizing it.

## DISCUSSION

We presented a simple, reproducible and effective seedling microbiota engineering method using SynCom inoculation on bean seeds. The method was successful using a wide diversity of SynCom compositions (13 SynComs) and strains (36 strains) that are representative of the common bean seed microbiota. First, this method enables the modulation of seed composition and community size, even in a coalescence context with the native seed microbiota (*i.e* unsterilized seeds). Then, this SynCom colonization was effective in a second coalescence event with unsterilized potting soil. SynComs contributed on average to 80% of the seedlings’ microbiota. We showed that the mass effect was the main driver of seedling microbiota colonization. Additionally, individual strains showed variable seed and seedling colonization capacities that mostly depended on their phylogenetic affiliation. Finally, through priority effects, the engineered seed microbiota modified the overall seedling and rhizosphere microbiota assemblies.

### The use of SynCom to study ecological processes during seed and seedling microbiota assembly

Microbiota characterization on individual seeds demonstrated their low carrying capacity, their high variability in terms of diversity and composition (Chesneau et al 2022, Simonin et al 2022). Therefore, it is difficult to establish causality between seed microbiota composition and seedling microbiota composition. Here, we showed that common bean seed can be colonized by contrasted SynCom concentrations and compositions. This SynCom inoculation method could be an interesting strategy for improving our understanding of seed to seedling microbiota transmission and the ecological processes involved in plant microbiota assembly (Vorholt et al 2017). Especially, our metabarcoding approach using *gyrB* gene enables the tracking of individual strains within different niches and in different plant species (Simonin et al 2023). This allows a better understanding and characterization of colonization of each SynCom member of plant tissues, which is usually not possible using 16S rRNA gene and clustering approaches (Armanhi et al 2021). Thus, this microbiota engineering method enables more accurate sources-sink analysis by deeply decreasing the unknown source fraction contributing to seedling microbiota assembly in native seed communities (Rochefort et al 2021). Contrary to our initial hypothesis (H1.1) the primary source of microorganisms for the seedling was not the potting soil but were the native seed microbiota and the SynCom. The seed microbiota is thus an important source of microorganisms for seedling microbiota assembly, as reported by previous studies (Moroenyane et al 2021, Johnston-Monje et al 2021).

Strong changes in community composition are observed during seed germination and seedling emergence (Barret et al 2015). These microbiota shifts are mainly described as a consequence of a deep modification in plant physiology that leads to the selection of specific microorganisms (Torres-Cortés 2018, Chesneau et al 2022). Indeed, during seed germination, diverse seed exudates are secreted in the surrounding soil which influences the microbial communities and form the spermosphere (Nelson et al 2018, Aziz et al 2021). Also, neutral events such as dispersion and mass effect are expected to play a role during seedling microbiota assembly but are less described. In this context, the method exposed here is interesting to decipher the relative importance of neutral and selective processes during seed and seedling microbial community assemblies. Indeed, by manipulating the concentration of inoculum and varying the composition and richness of multiple SynComs we were able to better characterize the importance of mass effect and selection processes during seedling emergence.

### Mass effect during seed and seedling colonization

In our study model (common bean), seeds are colonized on average by 10^2^ CFU per seed (Chesneau et al., 2022), which may explain why SynCom inoculated over 10^6^ CFU/mL had completely taken over the native seed-borne community. In a coalescence framework, it means that mass effect is more important than the priority effects that the native strains could have benefited from (Debray et al 2021). Overall, we showed that despite the low natural carrying capacity observed on seed, common bean seeds can be colonized by different SynCom biomasses which is interesting for both theoretical and applicative frameworks.

Seed microbiota contributions to seedling microbiota under natural conditions is very variable from one study to another (Rochefort et al 2021, Chesneau et al 2022, Walsh et al 2021, Johnston-Monje et al 2021). One reason is that in natural conditions, when seed meets the soil, it also meets the soil microbiota and a diversity of possible community coalescences. Rillig et al (2015) expose that one important parameter to predict the coalescence outcome is the mixing ratio of the two communities. In our case, we deliberately manipulated the inoculum concentration of the seed microbiota to vary this ratio. By doing so, we showed that mass effect was a key factor of the community coalescence outcome, as expected in hypothesis (H1.2). Indeed, SynCom contribution to seedling microbiota was correlated with inoculated seed community size. In the same way, strains relative abundance on seedlings were correlated with their relative abundance on seeds. Thus, it seems that a minimum abundance in seed is needed to be able to colonize seedlings. In the same idea, Darrasse et al (2007) showed that to effectively infect a seedling with *Xanthomonas citri* pv. *fuscans*, a minimum of 10^3^ CFU per bean seed was needed (Darrasse et al 2007). Arias et al (2020) also confirmed that a minimum biomass of *Xanthomonas vasicola* pv. *vasculorum* on seed was necessary to effectively colonize the plant.

To conclude, we show that mass effect is the main parameter of seed and seedling colonization (H1.2). Through mass effect, the seed microbiota has an advantage during seedling colonization compared to microorganisms from other environmental sources. This is a very promising result that shows that seedling microbiota can be modulated using a limited amount of inoculum on seed that is sufficient to outcompete soil microorganisms (Rocha et al 2021).

### Selection processes during seed and seedling colonization

SynComs inoculated at the same concentration (10^7^ CFU/mL) but of different compositions show different seed colonization capacities. At the strain level, we also observed variable seed colonization capacities. It means that selection processes also occurred during seed colonization. These differences between strains mainly depend on their phylogenetic affiliation. For instance, Enterobacteriaceae are significantly enriched in seeds. These variations in seed colonization may arise from differences in the adhesive capabilities of strains, which are influenced by the secretion of prominent adhesins (Espinosa-Urgel 2000, Duque et al 2013). The selection that occurred during seed colonization seems to be the main bottleneck of seedling colonization, as we previously showed that being abundant on seed was important to colonize seedling. From an applied point of view, it could be thus interesting to develop specific seed coatings for strains with low seed colonization capacities (Rocha et al 2019). These coatings could include binding molecules, prebiotic and specific nutrients to maintain strain of interest on seeds.

Even if generally, the relative abundance of a strain on seed can predict its relative abundance on seedling, we also show that strains have contrasting seedling colonization capacities supporting our hypothesis (H2.1). These differences between strains depend on their phylogenetic groups. Torres-Cortés et al. (2018) showed that bacteria having copiotrophic strategies with rapid growth have better seedling colonization capacities via competitive exclusion processes. Consistently with our results, they showed that Enterobacteriales and Pseudomonadales were enriched during seedling emergence. In particular, we show that an *Enterobacter cloacae* was dominating the control seedling microbiota. *Enterobacter cloacae* was detected on seed but at a low relative abundance (5%), has already been described as an obligatory plant endophyte in another study (Madmony et al 2005) and presents opportunistic characteristics with genes implied in colonization processes and copiotrophic strategy (Guérin et al 2020, P. Roberts et al 1992). These different observations suggest that redundant copiotrophic strategies of these strains increase their fitness during spermosphere formation and seedling emergence. Indeed, the multiple nutrients that are released during these events gives them clear advantages that select them.

Beyond intrinsic strain capacity to colonize seedling, we also show that biotic interactions are involved during seedling microbiota assembly (H2.2). For instance, *Frigoribacterium sp* (CFBP9039) and *Microbacterium sp* (CFBP9034) have a better seedling colonization in SynCom 8B and 11A than in SynCom 5C. Interestingly, their improved seedling colonization appears in SynCom with a higher richness level. This might highlight some facilitating processes (*e.g* niche expansion) from the other strains (Li et al 2018). On the contrary, *Sphingomonas sp* (CFBP9021) is never transmitted in SynCom 11A but has a transmission rate of 75% in SynCom 5A and 8B which might highlight some exclusion through interference (i.e. antagonism) or exploitative (*i.e*. niche occupation) competition (Hibbing et al 2010). Thus, strain colonization is SynCom dependent, as previously demonstrated in other plant habitats or ecosystems (Simonin et al 2023, Jones et al 2022). This confirms the complexity of biotic interactions during the seedling microbiota assembly and within SynComs and it entails considering these interactions when designing SynCom for microbiota engineering.

Overall, multiple selection processes also occurred during seed and seedling colonization. Strains showed variable seed and seedling colonization that depend on their phylogenetic affiliation and biotic interactions within the SynCom also influence the colonization success of the different strains. More studies are needed to elucidate the relative importance of host selection, environmental filtering and biotic interactions during seedling microbiota assembly. Also, further investigations are needed to better characterize strain transmission pathways in plant tissues and their stability during the plant development.

### Impact of seed microbiota on rhizosphere and seedling community assembly

We showed that SynComs inoculated on seed induce deep changes on the overall recruited communities from environmental sources (e.g soil, native seed community, H3.1). Seed microbiota is expected to highly interact with soil microbiota during the spermosphere assembly which in return influences the overall rhizosphere and plant microbiota assemblies (Aziz et al 2022, Olofintila & Noel 2023). Johnston-Monje et al (2016) showed that most seedling rhizosphere bacteria were seed derived. On the contrary, we showed that our SynCom strains contribution to the rhizosphere is low (< 0.2%) and identical to the control. Guo et al (2021) also showed that contribution of seed microbiota to the assembly of the rhizosphere microbiota was negligible (Guo et al 2021). In our case, even if the SynCom14 strain contribution was low, the overall rhizosphere community assembly was deeply modified (H3.2). This modification could arise from the priority effect of the inoculated strains on the initial spermosphere microbiota assembly (Aziz et al 2022) and subsequent developing rhizosphere community. Our results are consistent with Ridout et al (2019) showing a similar pattern with seed endophytes that influence the rhizosphere colonization of secondary symbionts through priority effect. The engineered seed communities could have changed the whole rhizosphere assembly through niche modification, biotic interactions and/or host control modification (Xu et al 2022). For instance, Kong et al (2021) showed that the inoculation of a plant using a specific *Bacillus amyloliquefaciens* strain could induce changes in volatile organic compounds (VOCs) emission that led to deep rhizosphere modification (Kong et al 2021). In the same way, co-inoculation of *Mesorhizobium ciceri* and *Bacillus subtilis* on seed induced changes in root exudates and rhizosphere microbiota assembly of chickpea (Scherbakov et al 2017). Rhizosphere assembly modification could also come from biotic interactions during the coalescence between the SynCom and the potting soil communities (Rocca et al 2021, Aziz et al 2022). Then, because the rhizosphere is one of the main sources of microorganisms for plant microbiota (Xiong et al 2021), it could be a factor explaining the differences observed in the recruited communities between the different SynComs. Indeed, we also showed that the extensive seedling microbiota assembly was deeply modified by the SynCom inoculation on seeds. In this context, we can argue that multiple successive selective processes led to the extensive rhizosphere and seedling microbiota assembly changes. At first glance, the SynCom can indeed colonize seedling and rhizosphere and interact with native communities. Then, through seedling physiological modification, the entire seedling and rhizosphere niches could be modified leading to differences in microbial colonization.

From a risk assessment point of view, the low colonization capacity of the inoculated seed microbiota into the rhizosphere could be taken as an advantage point. Indeed, the SynCom strains showed low environmental invasion capacities while still modifying the recruitment of native bacteria from the environment. Overall, our study shows that seed microbiota, through priority effect, is of great interest for microbiota engineering to modulate the overall seedling and rhizosphere microbiota assemblies.

## ACKNOWLEDGMENTS

This research was conducted through the OSMOSE project (2020-2022) in the framework of the regional programme “Objectif Végétal, Research, Education and Innovation in Pays de la Loire”, supported by the French Region Pays de la Loire, Angers Loire Métropole and the European Regional Development Fund. This work was also part of the 3^rd^ Programme for Future Investments (France2030) and operated by the SUCSEED project (ANR-20-PCPA-0009) funded by the ‘Growing and Protecting crops Differently” French Priority Research Program (PPR-CPA), part of the national investment plan operated by the French National Research Agency (ANR).

We thank Muriel Bahut (ANAN platform, SFR Quasav) for amplicon sequencing. The authors declare that they have no conflicts of interest.

---

**Figure S1:**
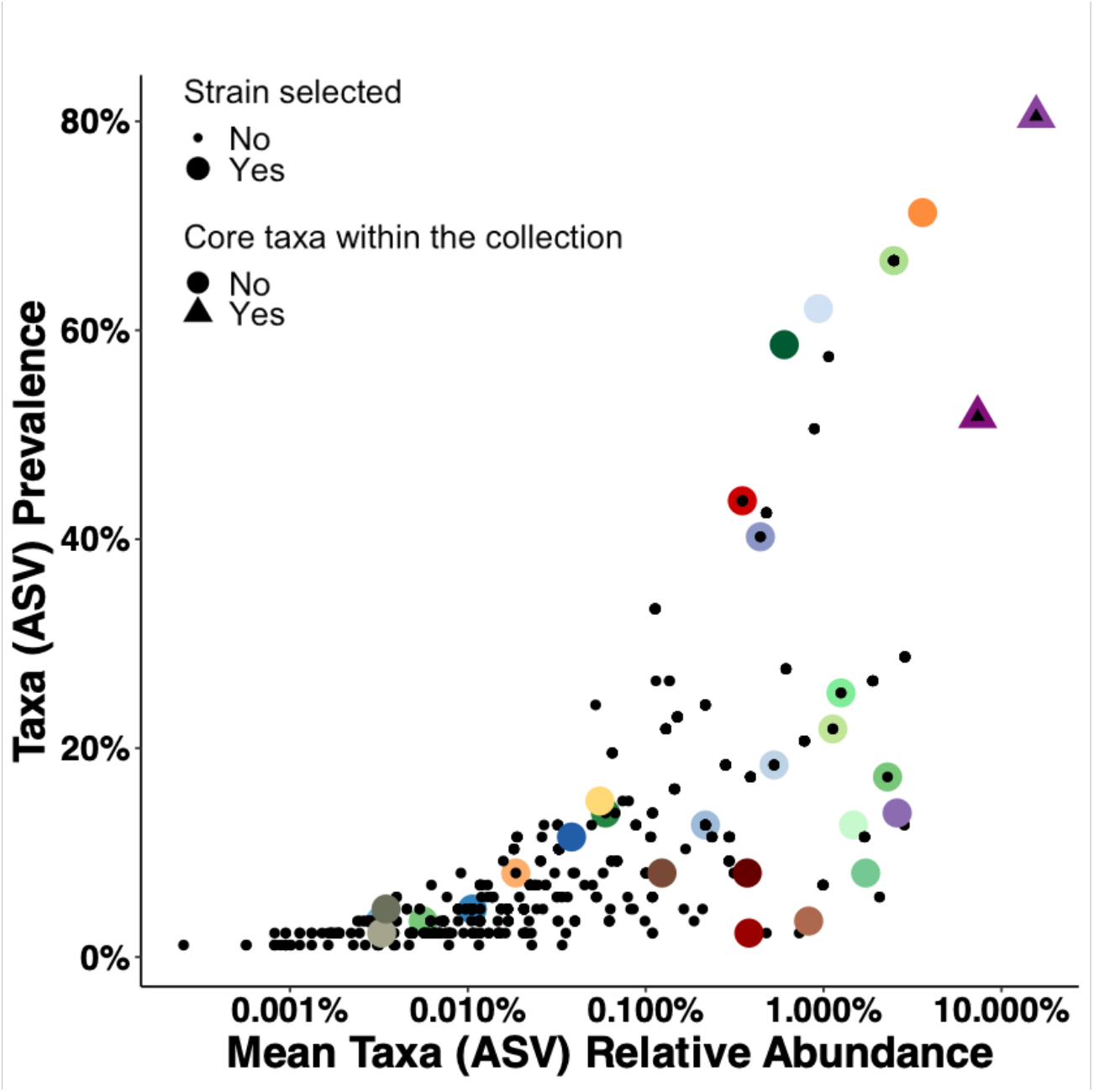
Strain selection for SynCom design. A total of 36 strains were selected (colored) within a collection of 1276 bacterial strains isolated from seeds and seedlings of 8 different genotypes of common bean. Strains were selected based on their prevalence and relative abundance in the common bean seed microbiota. Seven strains that were not detected in metabarcoding were also selected to include rare taxa.

**Figure S2:**
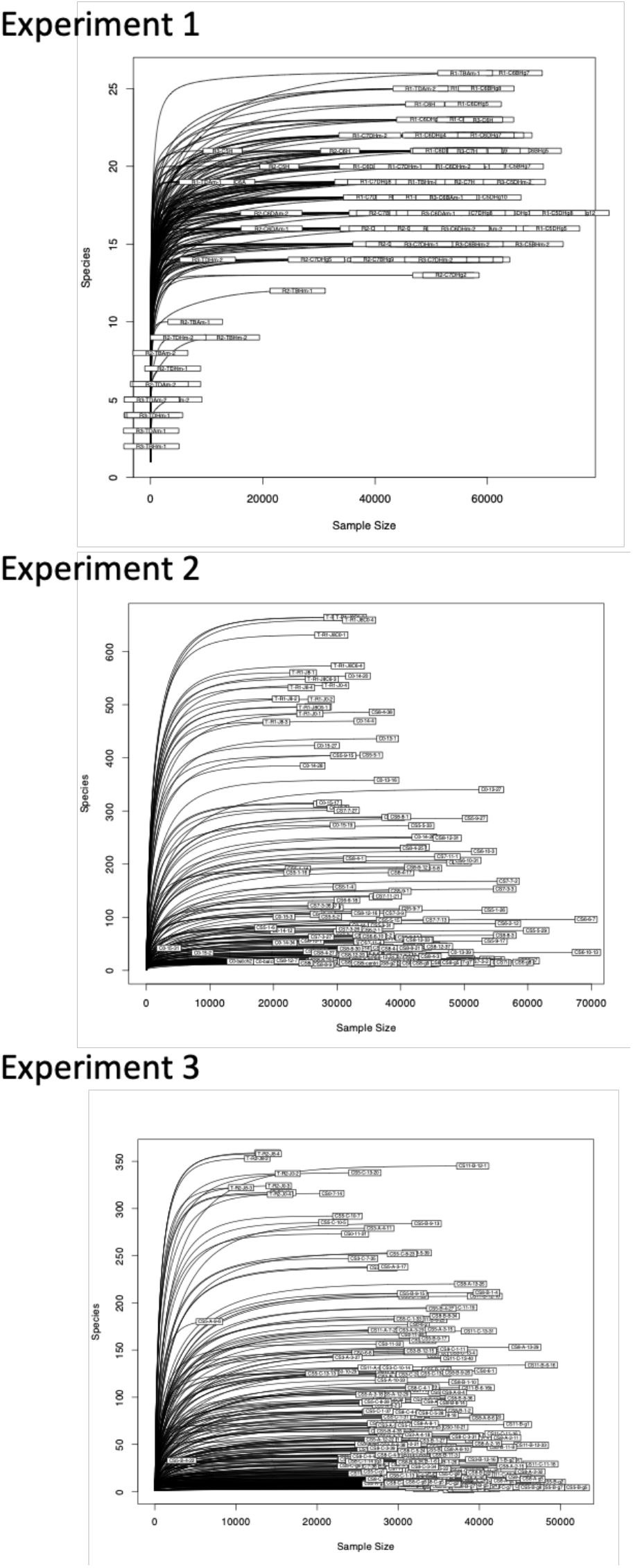
Rarefaction curves for the different experiments. Rarefaction curves for each sample of experiment 1, 2 and 3. For ordination analysis, datasets were rarefied at 10000 reads for experiment 1, 6000 reads for experiment 2 and 10000 reads for experiment 3. For alpha diversity analysis, datasets were rarefied at the minimum coverage from each experiment.

**Figure S3:**
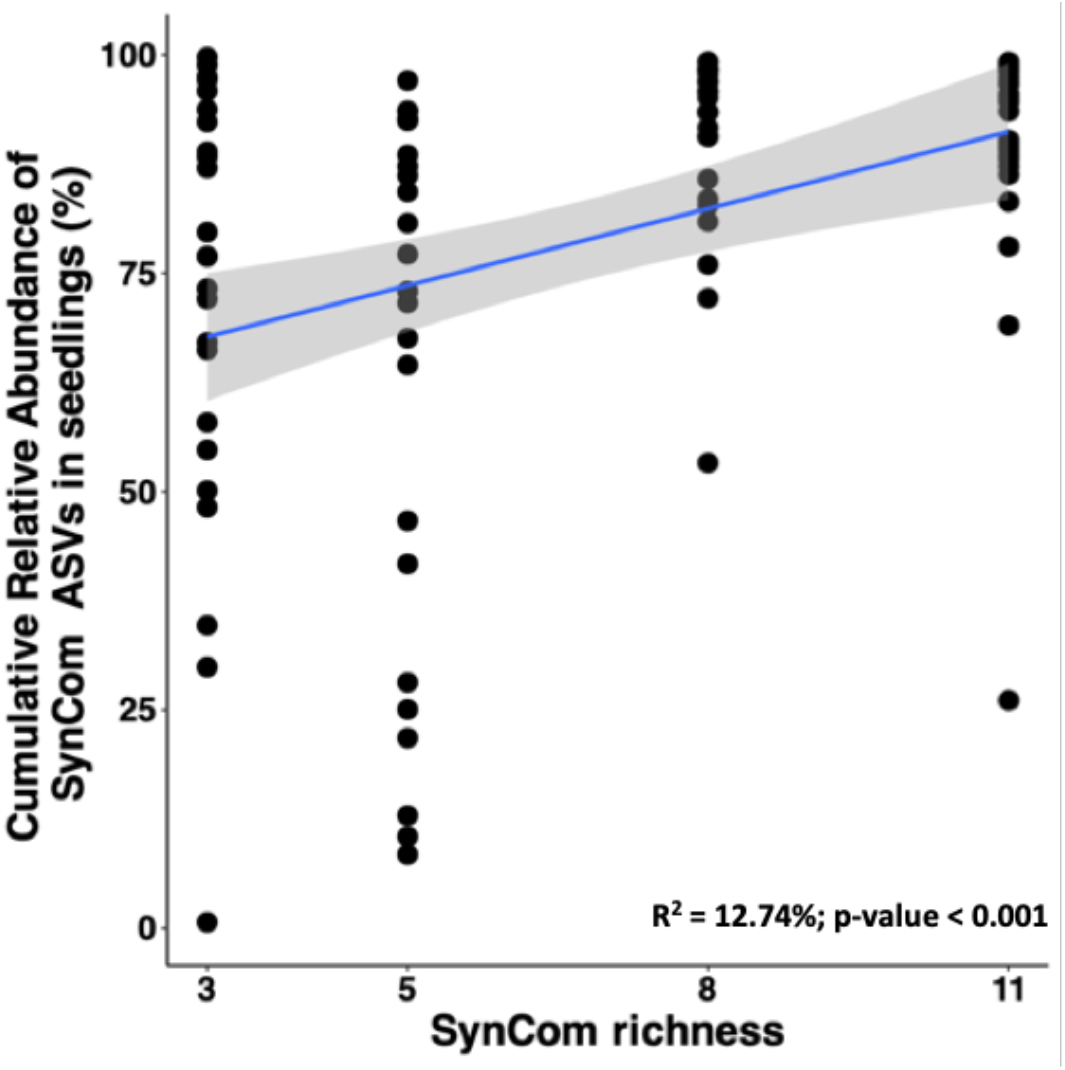
Correlation between cumulative relative abundance of SynCom ASVs in seedlings and SynCom richness.

**Figure S4:**
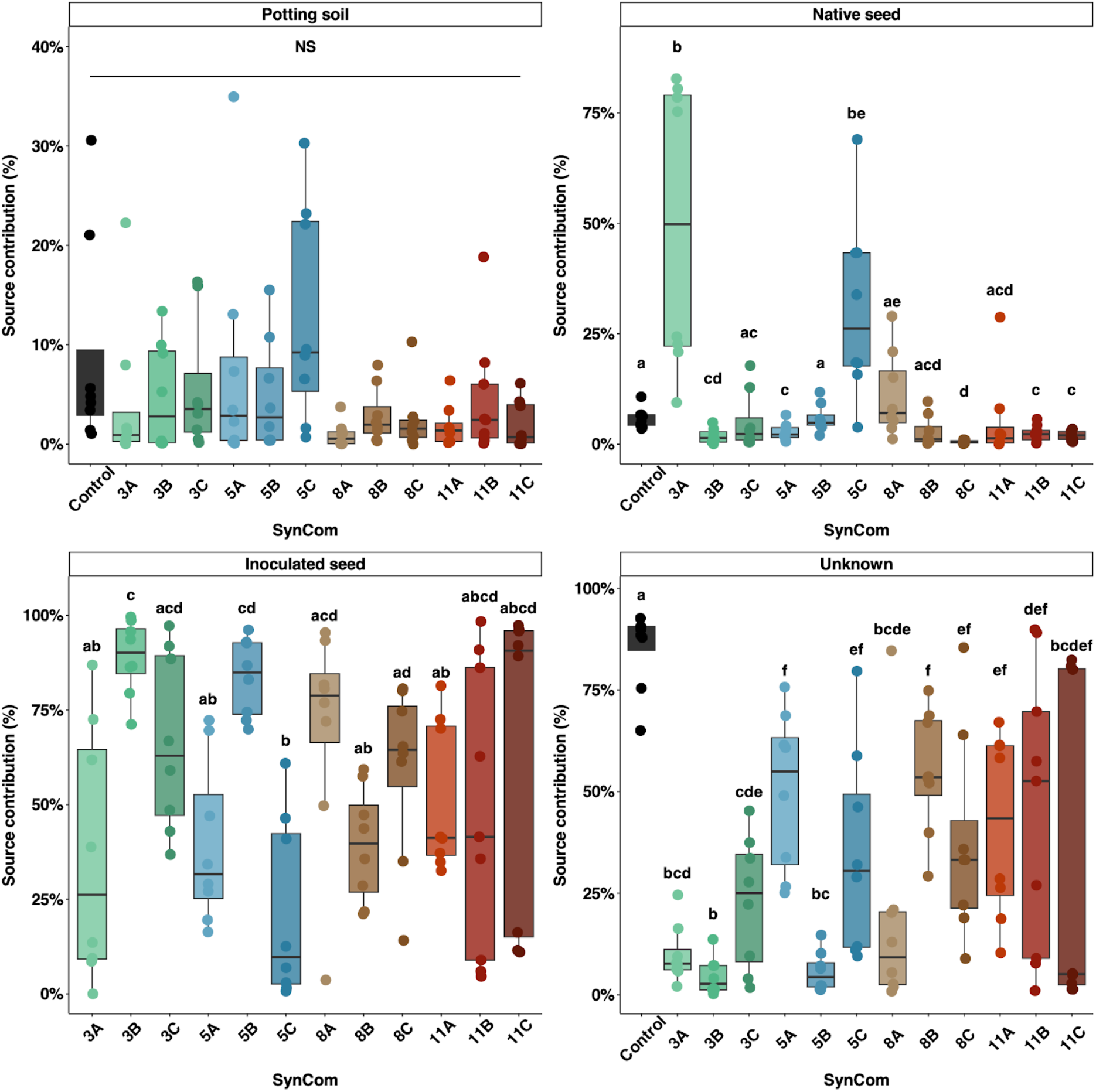
Relative contribution of native seed microbiota, potting soil and inoculated seed, using a microbial source tracking analysis (FEAST). Native seed, inoculated seed and potting soil microbiota were considered as sources of microorganisms and seedling were considered as sink. Detailed boxplot for each source and SynCom. The different letters indicate the significantly different groups (pairwise Wilcoxon, p-value < 0.05, corrected using Benjamini-Hochberg method).

**Figure S5:**
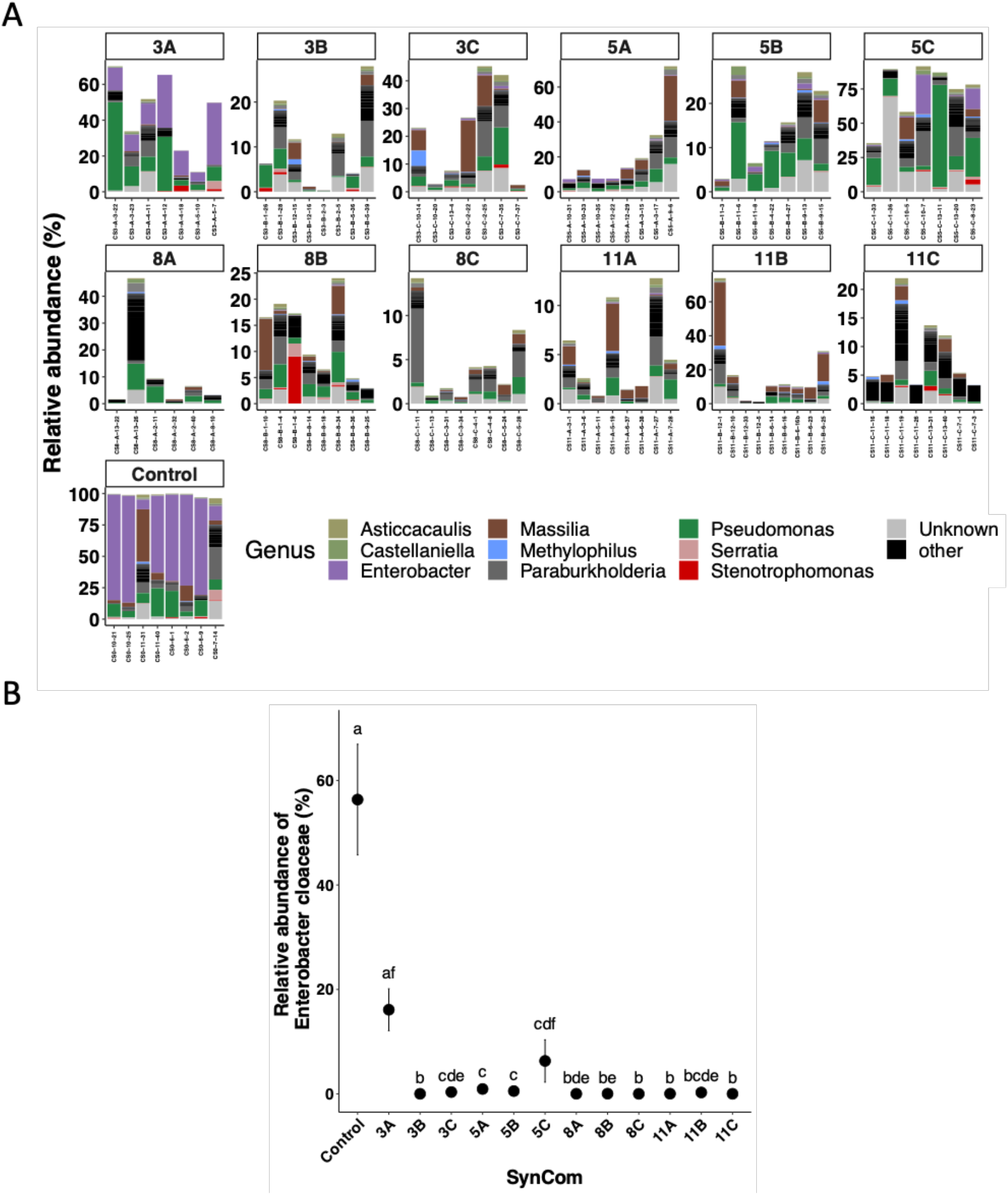
effect of SynCom composition on seedling recruited communities. A) Taxonomic profiles of the seedlings in the different SynComs of experiment 3. Each stacked bar represents a sample. ASVs of the SynCom strains were removed to plot only the recruited fraction from the environment and the native seeds. ASVs were agglomerated at the genus level and genus that represented less than 0.1% were put in the ‘other’ category (black). B) Focus on Enterobacter cloaceae relative abundance in seedlings of experiment 3 (the different letters indicate the statistically different groups (p-value < 0.05) using pairwise Wilcoxon test, p-value were corrected using Benjamini-Hochberg correction).

